# Benchmarking automated detection and classification approaches for monitoring of endangered species: a case study on gibbons from Cambodia

**DOI:** 10.1101/2024.08.17.608420

**Authors:** Dena J. Clink, Hope Cross-Jaya, Jinsung Kim, Abdul Hamid Ahmad, Moeurk Hong, Roeun Sala, Hélène Birot, Cain Agger, Thinh Tien Vu, Hoa Nguyen Thi, Thanh Nguyen Chi, Holger Klinck

**Affiliations:** K. Lisa Yang Center for Conservation Bioacoustics, Cornell Lab of Ornithology, Cornell University, Ithaca, NY, USA; Institute for Tropical Biology and Conservation, Universiti Malaysia Sabah (UMS), Kota Kinabalu, Sabah, Malaysia; Jahoo, Angdoung Kraleung Village, Sen Monorom Orang, MondulKiri, Mondulkiri Province, Cambodia; Wildlife Conservation Society, Cambodia, 21, Street 21 Sangkat Tonle Bassac, Phnom Penh, Cambodia; Department of Wildife, Vietnam National University of Forestry, Xuan Mai, Chuong My, Ha Noi, Vietnam; The Institute for Tropical Biodiversity and Forestry, Xuan Mai, Chuong My, Ha Noi, Vietnam; Bac Giang Agro-Forestry University, Bich Son Commune, Viet Yen District, Bac Giang Province, Vietnam

## Abstract

Recent advances in deep and transfer learning have revolutionized our ability for the automated detection and classification of acoustic signals from long-term recordings. Here, we provide a benchmark for the automated detection of southern yellow-cheeked crested gibbon (*Nomascus gabriellae*) calls collected using autonomous recording units (ARUs) in Andoung Kraleung Village, Cambodia. We compared the performance of support vector machines (SVMs), a quasi-DenseNet architecture (Koogu), transfer learning with pretrained convolutional neural network (ResNet50) models trained on the ‘ImageNet’ dataset, and transfer learning with embeddings from a global birdsong model (BirdNET) based on an EfficientNet architecture. We also investigated the impact of varying the number of training samples on the performance of these models. We found that BirdNET had superior performance with a smaller number of training samples, whereas Koogu and ResNet50 models only had acceptable performance with a larger number of training samples (>200 gibbon samples). Effective automated detection approaches are critical for monitoring endangered species, like gibbons. It is unclear how generalizable these results are for other signals, and future work on other vocal species will be informative. Code and data are publicly available for future benchmarking.

## 1. Introduction

### 1.1 Passive acoustic monitoring

The use of passive acoustic monitoring (PAM), a technique that utilizes autonomous acoustic recording units (ARUs), has seen a steady increase in terrestrial environments in recent years due to decreasing costs of the units along with increases storage capacity and battery life [1]. PAM can provide important insights on the abundance [2] and distribution of vocal animals across the landscape [3], environmental correlates of vocal behavior [4], and even individual identity within a population [5]. The use of archival PAM, wherein acoustic data are stored locally on secure digital (SD) memory cards, can result in the accumulation of terabytes to petabytes of data over the life of a PAM project. Manually annotating spectrograms is time and cost prohibitive, therefore necessitating automated analytical approaches [6].

### 1.2 Automated detection

Automated detection of acoustic signals is generally focused on determining the start and stop time of signal(s) of interest from longer recordings [7]. Early approaches for automated detection included spectrogram cross-correlation [8] and a combination of a band-limited energy detector with a traditional machine learning algorithm (e.g., support-vector machine) [9,10]. Recent advances in deep learning have led to substantial improvements in automated detection over traditional signal processing approaches [11]. Many of these applications have focused on avian species, with BirdNET [12] and Perch (https://github.com/google-research/perch; [13]), being two applications that were developed for bird call classification. BirdNET has since been used for automated detection of many different taxa, including Yucatán black howler monkey (*Alouatta pigra*) [14]; in this application howler monkey sounds were incorporated into the original model training data.

Another open source model, Koogu (https://github.com/shyamblast/Koogu), is based on a quasi-DenseNet architecture and was chosen for relative computational efficiency and comparatively high performance with a small number of training samples [15]. Koogu has been used successfully for automated detection applications for katydid *spp*. [16,17], fin whales (*Balaenoptera physalus*) [15], and gibbons [18]. Transfer learning allows for the use of embeddings from models trained in one domain or on a particular dataset for a new problem. Early applications of transfer learning for bioacoustics included using embeddings from models trained on the large-scale image databases for classification of spectrogram images [19]. Recent applications of transfer learning, wherein embeddings from BirdNET and Perch— models that were trained on large datasets of avian sounds—were used to train new classifiers, and had superior performance compared to other models evaluated for a variety of mammalian species including marine mammals and bats [20].

### 1.3 Gibbons

Due to their loud calls that can be heard by human observers over 1.5 km in the dense forest [21], gibbons are ideal candidates for PAM. Automated detection/classification of gibbon sounds has been done thus far for five gibbon species, Hainan gibbons (*Nomascus hainanus*) [22], western black-crested gibbon (*N. concolor*) [23], Northern grey gibbons (*Hylobates funereus*) [10,24], Bornean white-bearded gibbon (*H. albibarbis*)[18], and southern yellow-cheeked crested gibbon (*N. gabriellae*) [24]. Previous work investigating the performance of six convolutional neural network (CNN) architectures (AlexNet, VGG16, VGG19, ResNet18, ResNet50, ResNet152) pretrained on the ‘ImageNet’ dataset [25] for automated detection for southern yellow-cheeked crested gibbon calls (the species included in the present study) found that ResNet50 architecture had the highest performance [24]. As both PAM and automated detection approaches become more accessible, more gibbon species will be included.

### 1.4 Objectives

Here, we compare a traditional machine learning approach (Mel-frequency cepstral coefficients [26] plus support vector machines [27]), to a deep-learning approach (Koogu) [15] and two transfer learning approaches: the use of a ResNet50 [28] CNN pretrained on the ImageNet dataset [25] and BirdNET [12,20] for the automated detection of southern yellow-cheeked crested gibbon (*N. gabriellae*) calls. Our objectives include: 1) investigating the performance of binary or multi-class models for a binary classification task of gibbon versus noise; 2) testing the impact of varying number of training samples on the performance of the algorithms for automated detection; and 3) evaluate the performance of models for binary classification of calls from a closely related gibbon species to determine the generalizability [7]. For the multiclass models we included training samples from another gibbon species (*Hylobates funereus*), that are found in North Borneo [10]. We expected that given the recent promising results for using embeddings from this model for non-avian species [20], BirdNET would outperform the other algorithms for all tasks, exhibiting higher performance with a fewer number of training samples. The data are publicly available on Zenodo (https://zenodo.org/records/12706803) for future benchmarking work on gibbon automated detection.

## 2. Methods

### 2.1 Acoustic data collection

We collected acoustic data from March-September 2022 (training data) and August – November 2022 (test data) in the ecotourism site Jahoo, located in Andoung Kraleung Village within the Keo Seima Wildlife Sanctuary, Cambodia. We used SwiftOne autonomous recording units (K. Lisa Yang Center for Conservation Bioacoustics, Ithaca, NY, USA). For the training data, 10 units were placed (∼250 m spacing) in the territories of two gibbon groups. For the test data, 10 units were placed on a separate, non-overlapping array at ∼2 km spacing. The arrays used for training and testing were approximately 2 km apart. All units recorded continuously over the 24-hour period at a sample rate of 32 kHz and 32 dB gain. Sound files were saved as 1-hour waveform audio (.wav) files, with a file size of approximately 230 MB. See [24] for more details on acoustic data collection and Figure 1 for a map of the PAM arrays.

**Fig 1.**
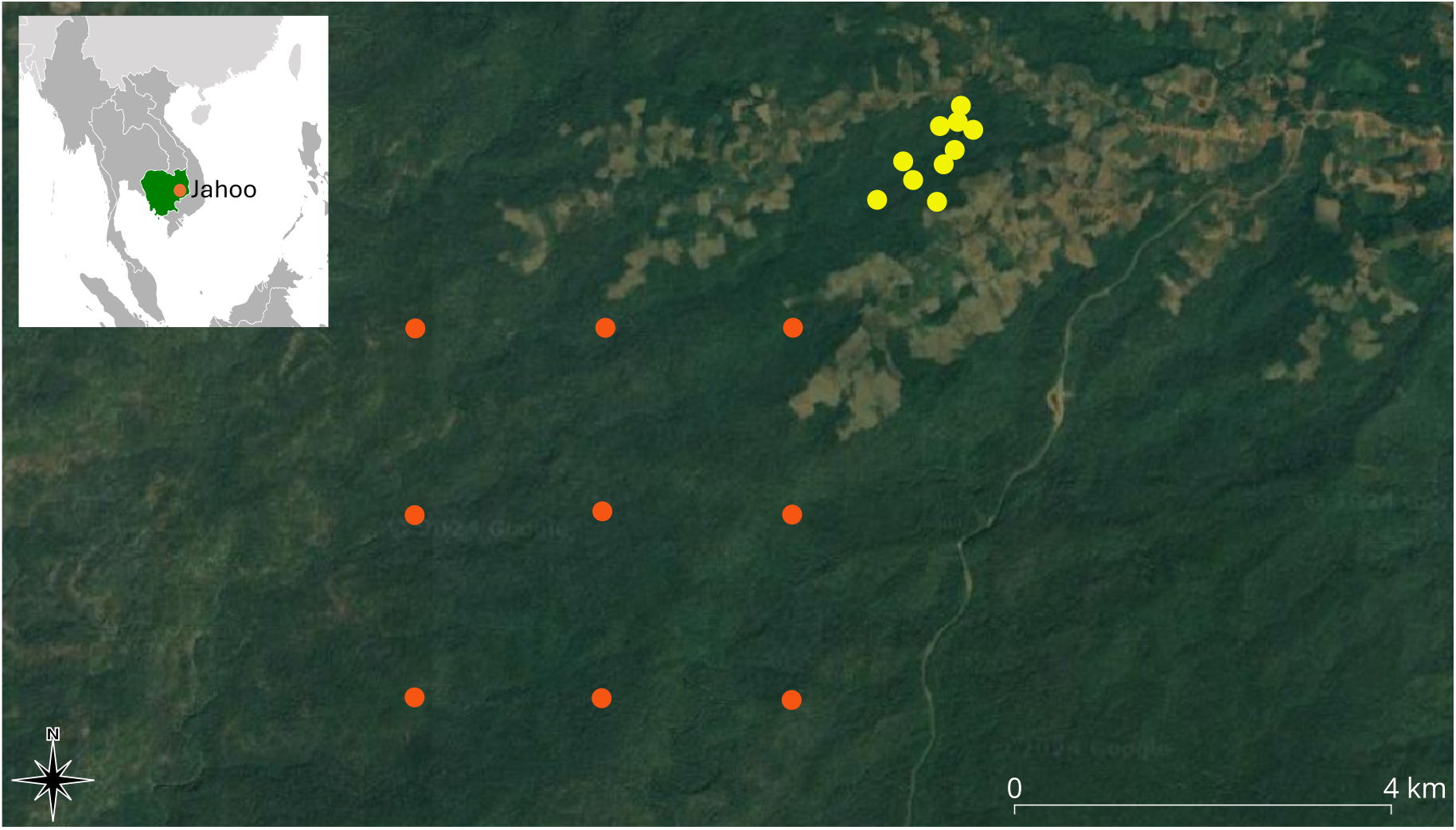
Map of PAM arrays in Jahoo in Andoung Kraleung Village, Cambodia used for training (yellow points) and testing (orange points; B). The inset map was modified from ASDFGHJ (Public Domain, https://commons.wikimedia.org/w/index.php?curid=7350825) and the larger map with the PAM points was made using QGIS v 3.34.1-Prizren (QGIS Geographic Information System. Open Source Geospatial Foundation Project. http://qgis.org).

## Training and test data preparation

To create the training dataset, two analysts (JK and HCJ) manually annotated 789 hours of randomly selected 1-hour recordings between the hours of 05:00 and 10:00 LT (as gibbons tend to call in the morning hours [29]) across the training array, recording all instances of female gibbon calls in spectrograms created in Raven Pro 1.6.3 (K. Lisa Yang Center for Conservation Bioacoustics, Ithaca, NY, USA, 2023). Raven settings were set to window size = 2,400 samples, contrast = 70, and brightness = 65; all other settings were the default. See Figure 2A for a representative spectrogram of a yellow-cheeked crested gibbon duet. We also indicated whether the calls were high-, medium- or low-quality, and omitted the low-quality calls from our training dataset; this resulted in 213 gibbons calls. We assigned calls to the ‘high-quality’ category if they had visible harmonics, indicating the calling animals were close to the recording unit. Whereas medium-quality calls had few visible harmonics but still had clear visible structure stereotypical of the gibbon female call. We considered calls ‘low-quality’ if they had low signal-to-noise ratio (<10 dB signal-to-noise ratio) or exhibited substantial overlap with another non-gibbon signal. We omitted low-quality calls from our training data (see Figure 2 B-D for representative spectrograms of call quality categories), and we combined ‘medium-’ and ‘high-’ quality calls for training. To create a noise class, we randomly selected 2,130 12-s clips from recordings that did not contain gibbon calls. The mean duration of the female gibbon call in this species is ∼ 12-s, with fundamental frequency range between 1 kHz – 3.5 kHz.

**Fig 2.**
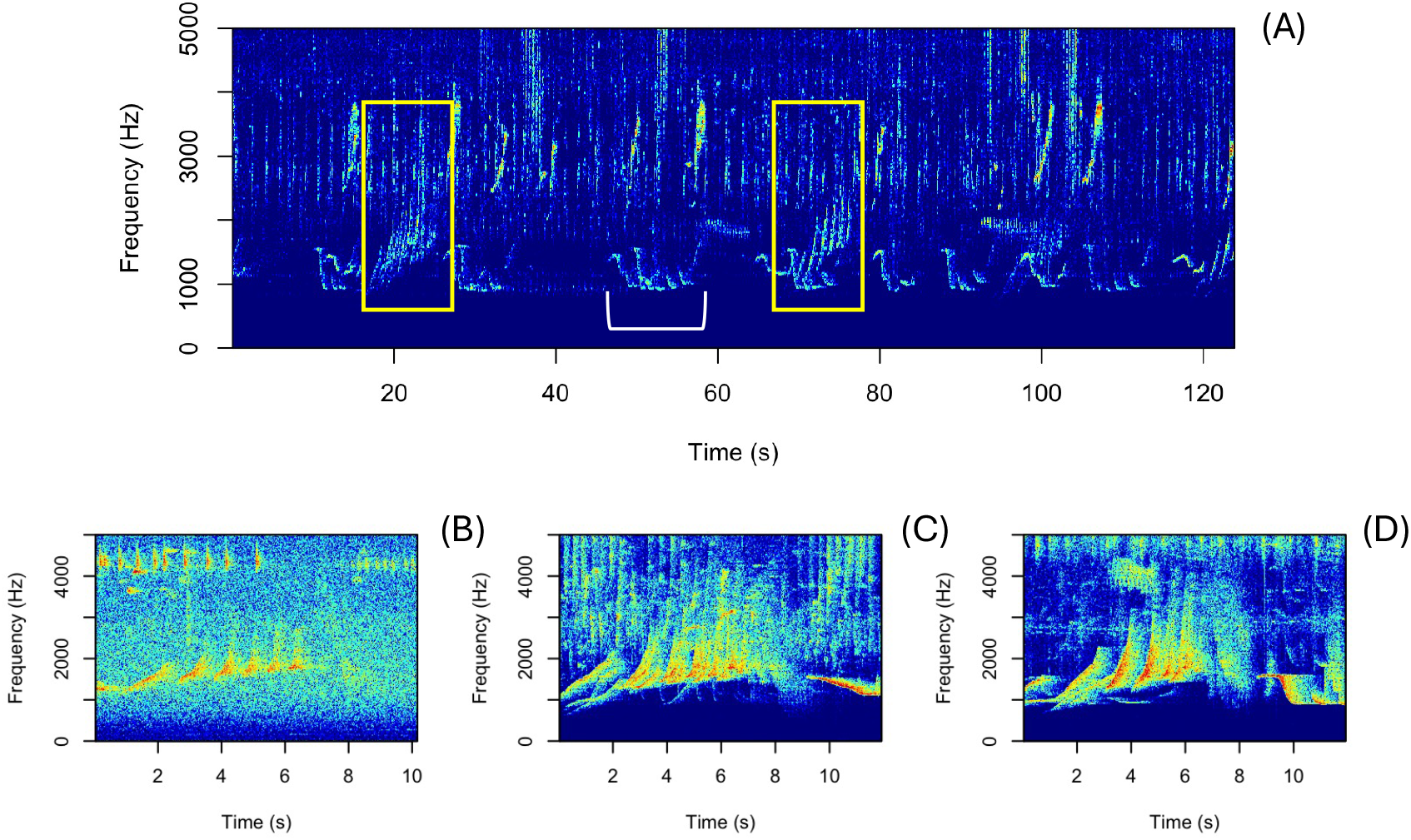
A representative spectrogram of a southern yellow-cheeked crested gibbon duet (A) and gibbon calls of varying quality (B-D). In plot A the yellow boxes indicate the female contribution to the duet, and the white bracket indicates the male contribution. For clarity, only one male contribution is indicated with the brackets. In plots B-D female calls of varying quality are shown, with B representing a “low-quality” call, C a “medium-quality” call, and D a “high-quality” call. For training, we omitted all “low-quality” calls.

We randomly selected 5, 10, 15, 20, 25, 45, 65, 85, 105, 125, 165, and 185 gibbon or noise samples over 5 iterations to examine the impact of number of training samples on model performance. We also included the full training dataset described above, along with a separate training dataset that was comprised of 1,090 gibbon samples and 1,748 noise samples that were automatically detected using early experiments with a binary AlexNet architecture [30] trained on the full dataset in the ‘torch for R’ ecosystem [31,32]. The detections were manually sorted by DJC into the gibbon or noise class by visual inspection of spectrogram images. For the multi-class models, we used an additional dataset of 502 Northern grey gibbon calls recorded in Danum Valley, Sabah, Malaysia. See [10,24] for details on data collection and preparation for this open dataset.

To create a test dataset, a single observer (DJC) manually annotated 10 hours of recordings confirmed to have gibbon calls present using Raven Pro 1.6; in this case, all default settings were used (window size = 512 samples, contrast = 50, and brightness = 50). As noted above, the test dataset came from a different PAM array to ensure that training and test datasets were independent [7]. Using the bounding boxes created using Raven Pro 1.6.3 selection tables, we created two separate formats for the test dataset. For the first format, which is more appropriately considered a classification problem, we divided the recordings into 12-s clips with 6-s overlap, and assigned any clips that contained at least 9-s of gibbon call annotations to the gibbon class, and the rest of the clips were classified as noise. A single observer (DJC) manually removed any clips in the noise category that contained partial gibbon calls. This resulted in 118 gibbon clips and 5,684 noise clips. The second format was more appropriate for ‘automated detection’, wherein we aimed to identify the start and stop time of gibbon calls within longer recordings. For this approach, we deployed the trained algorithms over the full recordings (see below for specifications) and had them output Raven selection tables. To evaluate model performance, we matched the selection table outputs to the start time of manual annotations. BirdNET returns predictions for each 3-s window, whereas ResNet50 returns predictions for 12-s windows, so we considered any detection within 6-s of the start and 3-s after the start of the annotation to be a detection. Consecutive detections were pooled so that there was only one detection per manual annotation. See below for more details.

## Assessing the generalizability of the models

To test the generalizability of the models, we used a different test dataset from Dakrong Nature Reserve in Vietnam that contains recordings of a recently recognized distinct gibbon species: the Northern buff-cheeked gibbons (*N. annamensis*). These gibbons were originally classified as *N. gabriellae* [33]. This dataset was comprised of 45 gibbon samples and 173 noise samples at a 16 kHz sample rate. BirdNET automatically up-samples all training and test sounds to 48 kHz, and for ResNet50 the input to the model is an image so it is not necessarily impacted by varying sample rates, although different sample rates do influence spectrogram resolution when using the same settings. See [24] for details and Table 1 for a summary of training and test data used in the present study.

**Table 1.** Summary of training and test datasets used in the current study.

| Dataset | Country | Species | Number of<br>Gibbon<br>samples | Number of<br>Noise<br>samples | Sample<br>rate | Data<br>format |
| --- | --- | --- | --- | --- | --- | --- |
| Training data |  |  |  |  |  |  |
| Jahoo Training<br>Data<br>(annotations) | Cambodia | Crested<br>gibbon | 213 | 2,130 | 32 kHz | Clips<br>obtained<br>using<br>manual<br>annotations |
| Jahoo Training<br>Data (sorted<br>detections) | Cambodia | Crested<br>gibbon | 1,085 | 1,748 | 32 kHz | Sorted<br>true/false<br>positive<br>clips from<br>early<br>experiments |
| Danum Valley<br>Conservation<br>Training Data | Malaysia | Grey<br>gibbon | 502 | ~ | 16 kHz | Clips, only<br>the positive<br>gibbon<br>samples used from<br>this dataset |
| Test data |  |  |  |  |  |  |
| Jahoo Test Data<br>(Clips) | Cambodia | Crested<br>gibbon | 118 | 5,684 | 32 kHz | 12-s clips<br>from sliding<br>time<br>windows |
| Jahoo Test Data<br>(1-hr files) | Cambodia | Crested<br>gibbon | 111 | ~6,000 | 32 kHz | Models<br>were<br>deployed<br>over full 1-hr files |
| Dakrong Nature<br>Reserve Test<br>Data | Vietnam | Buff<br>cheeked<br>gibbon | 53 | 175 | 16 kHz | Clips from<br>manual<br>annotations |

### 2.2 Model training and deployment

#### 2.2.1 Support vector machine

We used the R package “tuneR” [34] to calculate Mel-frequency cepstral coefficients (MFCCs) for each labeled sound event. We calculated MFCCs focusing on the fundamental frequency range of female gibbon calls (0.5–3.0 kHz), as harmonics are typically not visible in the recordings unless the animals are in close proximity to the recording units. Due to the variable duration of sound events, and the necessity for feature vectors of equal length in machine learning classification, we created 9 equally spaced time-windows for each clip. We then calculated 12 MFCCs along with the delta coefficients for each of the 9 time-windows [34]. We appended the duration of each call to the MFCC vector, resulting in a feature vector of length 177 for each sound event. We trained a support vector machine (SVM) using the ‘e1071’ package [35] using a 25-fold cross-validation (cross = 25) and a radial kernel.

#### 2.2.2 Koogu

We used Koogu v0.7.2 [36] that relies on the TensorFlow [37] deep learning framework. Koogu takes spectrograms as inputs, and we used the ‘data settings’ function to specify the clip length (12-s) and sample rate (32 kHz) of input clips. We set the spectrogram settings as window length = 0.128 seconds with 75% overlap and a frequency range of 500 to 3,000 Hz. For model training, we set aside 15% of the data for validation. We set the batch size to 64 and allowed the model to run for 50 epochs. We followed the default learning rate set by Koogu developers, and set the learning rate to start at 0.01 and reduced it to 0.001 at epochs 20 and 40. The dropout rate was set to 0.05. These were all default settings for the Koogu Python package [36]. We used the ‘train’ function to train the model and the ‘recognize’ function to classify the clips.

#### 2.2.3 ResNet50 CNNs

Our previous work that compared multiple CNN architectures found that ResNet50 led to the highest performance [24], so we used only this architecture for our analyses. We used the ‘spectro’ function in the ‘seewave’ R package [38] to generate spectrogram images to input into the model. We used the default window size and color scheme, removed all axis labels, and specified a frequency range of 0.5 –3.0 kHz for crested gibbons. We saved spectrogram images as ‘.jpg’ files using the ‘jpeg’ function in base R, as this is the most common format for these image classification models.

At the model input stage, we converted the images to tensors, resized them to 224×224 pixels, and normalized them by mean and standard deviation using the ‘torchvision’ R package [32]. Previous studies have shown that fine-tuning the feature extractor layer along with the output layer improves performance [19]. Therefore, for all models, we set the ‘requires_grad_’ parameter to ‘TRUE,’ unfreezing the extractor layers. All models were pre-trained on the ‘ImageNet’ dataset [25] and are accessible via the ‘torchvision’ R package v 0.5.1 [32].

We assessed the model performance over 5 epochs, as this was shown to lead to the best performance on this dataset in previous work [24]. We set the maximum learning rate to 0.001 [39], the batch size to 32, and utilized the Adam optimizer [40]. We employed the one-cycle learning rate strategy [41] implemented via the ‘luz_callback_lr_scheduler’ function in the ‘luz’ R package [42]. We used the ‘nn_bce_with_logits_loss’ function for binary classification models, which combines a sigmoid layer with the binary cross entropy function. For multiclass models, we used ‘nn_cross_entropy_loss,’ which calculates the cross-entropy loss between inputs and targets and is suitable for multi-class classification tasks [31]. To evaluate model performance for the binary models, the output layer had a single dimension, so we used a sigmoid function to convert the output of the CNN predictions on the test datasets to values between 0 and 1. For the multiclass models, the output layer had three dimensions representing the three classes in our data, and we computed class probabilities by applying a softmax function. We used the ‘gibbonNetR’ package (v1.0.0-beta.1) for this analysis [43].

#### 2.2.4 BirdNET

Recent advances in transfer learning using BirdNET [20], means that we could use the feature embeddings from BirdNET (version V2.4) trained on vocalizations of over 6,000 avian species to train a new classifier. BirdNET V2.4 uses an EfficientNetB0-like backbone [44] and is trained predominantly on data from Xeno-canto (https://xeno-canto.org/) and Macaulay Library (https://www.macaulaylibrary.org/). We used the ‘train.py’ function implemented in BirdNET (https://github.com/kahst/BirdNET-Analyzer) for each training dataset. This function allows users to train a new classifier for classes not currently in BirdNET using BirdNET embeddings. To create predictions from the trained models, we used the ‘analyze.py’ function. This version of BirdNET returns predictions for 3-sec windows. For both training and analysis, we used the default settings apart from setting fmin = 500 Hz and fmax = 3,000 Hz. For the default settings, the number of training epochs was 50, the batch size was 32, the validation split ratio was 0.2 and the learning rate was 0.001. We utilized a single-layer classifier with no hidden units and did not apply dropout or mixup. The model outputs were saved in the ‘tflite’ format.

### 2.3 Model evaluation

We used two separate approaches to evaluate model performance. For the classification problem, we deployed models trained on all the high-quality training samples (n = 213 gibbon calls) over a test dataset consisting of 118 gibbon calls and 5,684 noise samples. The SVM, Koogu, and the pre-trained CNNs output a single prediction and associated probability for each clip, whereas BirdNET outputs a prediction for each 3-s window. For the BirdNET output, we identified the 3-s window within the 12-s clip with the highest confidence score and assigned the score and associated label to the clip.

We were interested to see how the number of training samples would influence the performance of our models, so we randomly selected 5, 10, 15, 20, 25, 45, 65, 85,105, 125, 149, 165, 185 gibbon samples and noise samples over five iterations. We also trained the models on 213 high-quality samples and 1,085 manually sorted detections from previous experiments. We then deployed the newly trained classifier over the test datasets and reported F1, precision/ recall using the ‘caret’ package (Kuhn, 2008), and area under the receiver-operating (AUC-ROC) curve calculated using the ‘pROC’ package (Xavier et al., 2011). For the automated detection approach, we used the manual annotations created in Raven Pro, and considered any detection within 6-sec of the start and 3-sec after the start of the annotation to be a detection.

### 2.4 Data and code availability

All code needed to recreate the analyses is on GitHub: https://github.com/DenaJGibbon/BirdNET-Performance-Comparison. Sound files are on Zenodo: https://zenodo.org/records/12706803.

## 3. Results

### 3.1 Evaluating binary and multi-class model performance using all training samples

We deployed binary and multi-class models (trained on 213 gibbon calls and 2,130 noise samples) over a test dataset comprised of 118 gibbon and 5,684 noise clips. We found that for this classification problem, the pre-trained ResNet50 CNN had a slightly higher F1 score than BirdNET (0.83 vs 0.71 for the multi-class models). Koogu had a slightly lower performance than the transfer learning approaches. For both the pretrained ResNet50 CNN and BirdNET, the multi-class models had better performance. We found that SVM + MFCCs led to very poor performance for this task (Figure 3).

**Figure 3.**
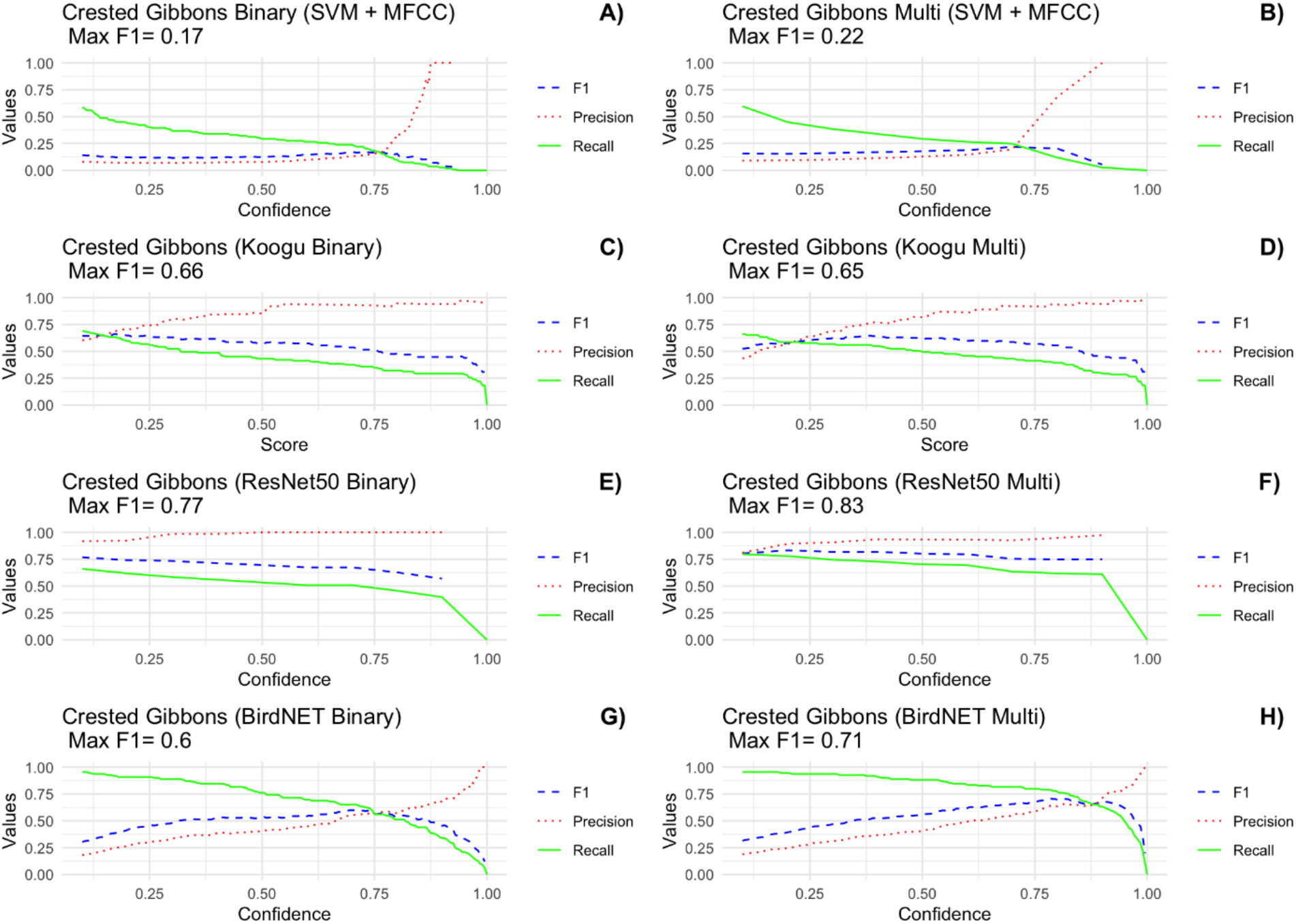
Precision, recall, and F1 score as a function of confidence for all models included in this benchmark on the Jahoo Test Data (clips). Models were trained on the full high-quality dataset, which contains 213 gibbon samples and 2,130 noise samples. See text for details on model configurations.

### 3.2 Evaluating model performance when varying number of training samples

We evaluated how varying the number of training samples impacted our results for an automated detection problem. We focused on ResNet50 CNN and BirdNET for this task, as the SVM + MFCC had very poor performance, and earlier experiments with Koogu showed that the model did not converge for tasks with very small training datasets. We found that BirdNET outperformed the ResNet CNN architecture at a lower sample size for both F1 and AUC-ROC scores (Figure 4). At a lower number of training samples, the ResNet50 CNN performed poorly, as indicated by a low F1 score and an AUC-ROC score worse than random chance (< 0.5). However, ResNet50 had a similar performance as BirdNET for both metrics with a higher number of training samples.

**Figure 4.**
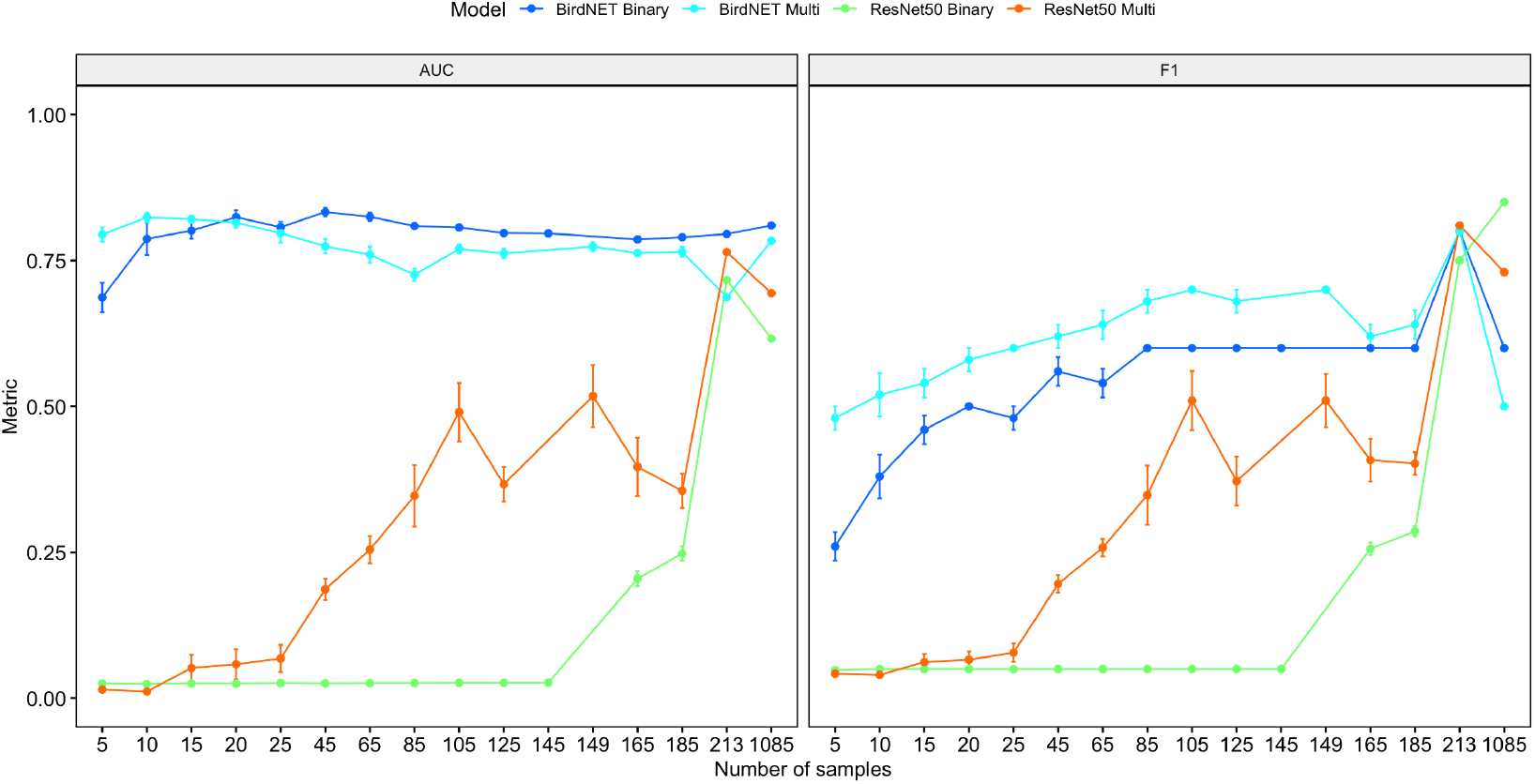
Performance as a function of the number of training samples for BirdNET and ResNet50 (CNN) over 5 iterations for an automated detection task on the Jahoo Test Data (1-hr files). In the plots above, the mean +/-standard error for the AUC-ROC score (A) and the maximum F1 score (B) are shown as a function of training samples for binary and multi-class models. For the number of training samples indicated by 213 and 1,085, there were no random iterations as all the training samples were used. An AUC-ROC value less than 0.5 indicates that the model performs worse than random chance.

### 3.3 Evaluating the generalizability of models trained on varying numbers of samples

To test the generalizability of the models, we evaluated the performance on a test dataset from a different location and gibbon species. We found that BirdNET outperformed the ResNet50 models (Figure 5), and for both architectures, the multiclass models outperformed the binary models in the majority of cases.

**Figure 5.**
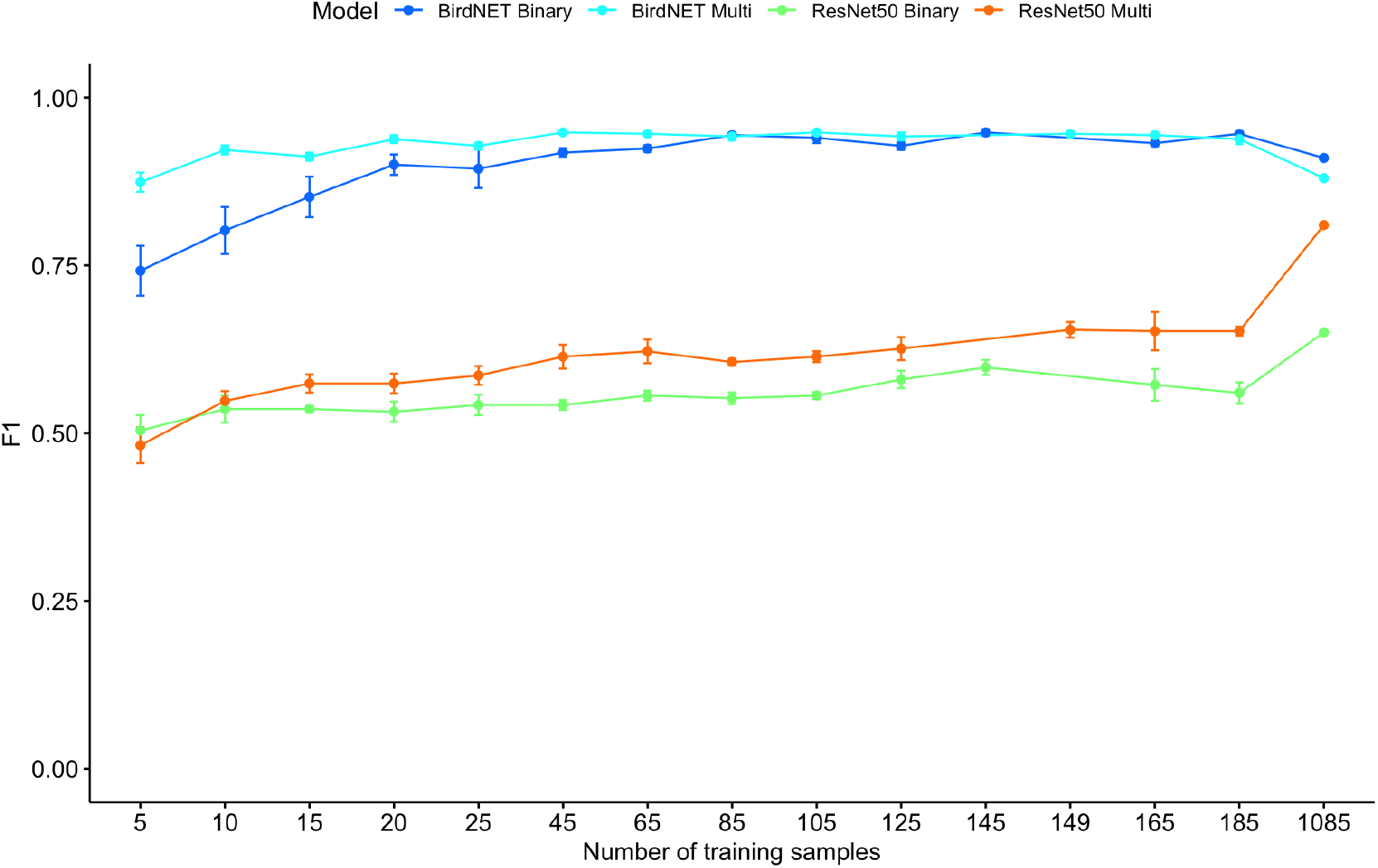
Performance as a function of number of training samples for BirdNET and ResNet50 over 5 random iterations for a classification task of calls from a closely related gibbon species (Buff cheeked gibbon) in Vietnam. The plots show the mean +/-standard error for the maximum F1 score for each model and training data combination. The training dataset with 1085 did not have random iterations, as the entire dataset was used for training. This analysis is meant to show the generalizability of the models.

## 4. Discussion

Here, we provide a benchmark for automated detection of gibbon calls from PAM data in Cambodia. We found that embeddings from BirdNET led to better performance with fewer samples but that the performance of ResNet50 models was comparable when the number of training samples increased. Koogu models had slightly lower performance on the full training dataset. Due to the very poor performance of ResNet50 with a small number of training samples, and the difficulties with a lack of convergence for Koogu models with smaller training datasets, we do not recommend these approaches for use with small training datasets.

In previous work on northern grey gibbons, the traditional machine learning approach (SVM + MFCC) combination leads to satisfactory performance for both automated detection [10] and classification of individual gibbons [45]. In addition, previous work found that for gibbon individual classification, the use of MFCCs and BirdNET embeddings combined with a traditional machine-learning approach (random forest) led to similar performance [46]. However, this was not the case with our dataset of yellow-cheeked crested gibbons. Our findings provide evidence that the performance of both the feature extraction method and the algorithm are highly dependent on the signal(s) of interest, even for closely related gibbon species. We believe these differences are probably due to the substantially different structure of the calls in the two different species.

Previous work using Koogu found a maximum F1 score of 0.87 for automated detection of Bornean white-bearded gibbon calls, whereas we found a maximum F1 score 0.66 for yellow-cheeked crested gibbons. There are a few possible explanations for the differences. First, we had fewer training samples, and did not perform data augmentation. It is possible that these additional steps would further improve model performance. The analyses were also done on different gibbon species, and it has been shown that the best-performing algorithm can vary depending on the gibbon species [24]. We opted not to use Koogu for the additional experiments varying the number of training samples as models did not converge after many epochs (>100) with a small sample size, and there was not an option for early stopping that would allow us to systematically evaluate model performance over many random iterations. In addition, Koogu model training on an Apple M2 Max with 12-core CPU, 30-core GPU, 16-core Neural Engine 64 GB unified memory took substantially longer for Koogu than the ResNet50 or BirdNET models. For example, using default Koogu settings (over 50 epochs) on the full training dataset it took more than one hour, whereas for BirdNET it took less than five minutes to train for 50 epochs on the same training dataset. Therefore, the difficulty with model training over multiple iterations and substantial training time meant that Koogu was not effective for our sample-size experiments.

Previous work on other bioacoustics signals found acceptable performance using transfer learning with models trained on the ImageNet dataset with a small number of training samples (<25 [19]). We found that model performance was highly dependent on the test dataset. For the automated detection problem, we found very low performance with smaller training datasets for ResNet50, but for the test dataset from Vietnam, we found better performance with a smaller number of samples. These differences highlight the importance of effectively curating test datasets to match the research question.

Our results provide evidence that embeddings from bioacoustics models, like BirdNET, lead to much better performance than other approaches for classification and automated detection of non-avian species [20]. Given the current biodiversity crisis, monitoring endangered species such as gibbons effectively is critical for effective conservation and management. We show that BirdNET can be used as an effective tool for the automated detection of southern yellow-cheeked crested gibbon calls from PAM data in Cambodia. Future work that benchmarks BirdNET for other gibbon species will be informative.

## Acknowledgments

We thank the Royal Government of Cambodia and the KSWS REDD+ program for funding fieldwork and Kyle Burrichter for his assistance with the early stages of field data collection at Jahoo.

## Author Declarations

### Ethics Approval

The research adhered to all applicable international laws, and it was conducted in Jahoo, Cambodia, with permission from the Royal Government of Cambodia.

